# Individual differences in the expression and control of anger are encoded in the same fronto-temporal GM-WM network

**DOI:** 10.1101/2024.05.29.596455

**Authors:** Alessandro Grecucci, Francesca Geraci, Ellyson Munari, Xiaoping Yi, Gerardo Salvato, Irene Messina

## Abstract

Anger can be deconstructed into distinct components: a temporary emotional state (state anger), a stable personality trait (trait anger), a tendency to outwardly express it (anger-out), or to internally suppress it (anger-in), and the capability to manage it (anger control). These aspects exhibit individual differences that vary across a continuum. Notably, the capacity to express and control anger are of great importance to modulate our reactions in interpersonal situations. The aim of this study was to test the hypothesis that anger expression and control are negatively correlated and that both can be decoded by the same patterns of grey and white matter features of a fronto-temporal brain network. To this aim, a data fusion unsupervised machine learning technique, known as transposed Independent Vector Analysis (tIVA), was used to decompose the brain into covarying GM-WM networks and then backward regression was used to predict both anger expression and control from a sample of 212 healthy subjects. Confirming our hypothesis, results showed that anger control and anger expression are negatively correlated, the more individuals control anger, the less they externalize it. At the neural level, individual differences in anger expression and control can be predicted by the same GM-WM network. As expected, this network included fronto-temporal regions as well as the cingulate, the insula and the precuneus. The higher the concentration of GM-WM in this brain network, the higher level of externalization of anger, and the lower the anger control. These results expand previous findings regarding the neural bases of anger by showing that individual differences in anger control and expression can be predicted by morphometric features.

## Introduction

Anger is a primary emotion typically characterized by discomfort serving to mobilize resources and enact change in response to provocation, hurt, or threat (Videbeck, 2006; Sorella et al., 2021; Sorella et al. 2022). Its constructive role encompasses facilitating goal achievements, obstacles overcoming, and maintaining interpersonal boundaries (Grecucci, Giorgetta, Bonini, & Sanfey, 2013; Grecucci, Giorgetta, van Wout, et al., 2013; Sorella et al., 2021; Sorella et al. 2022). Nonetheless, anger regulation is challenging due to the intense physiological reactions associated with the fight-or-flight response, activated to safeguard oneself from the provoking circumstances (Lazarus, 1991). Difficulties in regulating anger may serve as precursors of aggressive behaviours (Lochman et al., 2010), with consequent interpersonal difficulties and social maladaptation (Baron et al., 2006; Heilbron & Prinstein, 2008). And anger dysregulation is a core feature of psychiatric disorders such as borderline personality disorder (De Panfilis et al., 2019), antisocial personality disorder (Kolla et al., 2016), and intermittent explosive disorder (Coccaro et al., 2014).). Therefore, a comprehensive understanding of anger regulation is imperative. A widely recognized psychological framework aimed at elucidating the nature and components of anger regulation has been proposed by Spielberger and operationalized in the State-Trait Anger Expression Inventory (STAXI-2; Spielberger, 1999). This model recognizes anger expression and anger control as two main dimensions of anger regulation. Anger externalization (Anger Expression – Out) refers to an individual’s propensity to externalize or openly express their anger, through questions involving behaviours, actions, or reactions in anger-inducing situations, in contrast to anger internalization (Anger Expression – In) that regards the tendency of individuals to suppress or internalize anger. Anger control refers to people’s ability to control the physical or verbal expressions of anger (Anger Control-In) and to relax, calm down, and reduce angry feelings before they get out of control (Anger Control – Out). Among such dimensions of anger regulation, externalization and control represent complementary facets. Indeed, anger control can be viewed as the ability to restrain anger expression through control and outcome monitoring (Wilkowski & Robinson, 2008; Wilkowski & Robinson, 2010). If on the one hand externalizing anger in same cases may play a positive role (e.g., maintenance of boundaries) (Grecucci, Giorgetta, Bonini, & Sanfey, 2013; Grecucci, Giorgetta, van Wout, et al., 2013; Sorella et al., 2021; Sorella et al. 2022), on the other hand, when not counterbalanced by adequate anger control may have negative consequences. For instance, the combination between anger externalization and low anger control garners attention because both are considered precursors of hostility (Bridewell, 1997) and related social interaction issues (Baron et al., 2006) and aggressive behaviours (Roberton, 2015; Lochman et al., 2010; Birkley & Eckhardt, 2015). Similarly, it has been shown that people with poor control of anger have a greater propensity to angry externalization, fostering aggressive behaviours (Bettencourt et al., 2006; Dodge & Coie, 1987; Mattevi et al., 2019). From a clinical perspective, these dimensions of anger regulation require tailored interventions focused on externalizing psychological problems (distinct from those targeting internalized anger expression, more typical of internalizing problems) (Achenbach et al., 2016; Pascual-Leone et al., 2013; Manfredi & Taglietti, 2023).

Despite the importance of individual variances in regulating anger for mental well-being, there exists limited evidence regarding the neural foundations of these individual differences. At a structural level, Sorella et al. (2021) found that the concentration of grey matter in a network comprising ventromedial temporal areas, posterior cingulate, fusiform gyrus, and cerebellum correlated with trait anger. In line with such results, previous functional studies have found altered resting state activity (Sorella et al., 2021) and functional connectivity (Fulwiler et al., 2012) in the Default Mode Network (DMN). Beside such recent attempts to study the neural correlates of individual differences in trait anger, there is poor evidence on the neural bases of individual differences in anger regulation, neither in terms of anger expression nor anger control.

Related to externalization, some studies tried to understand the neural bases of aggression, a dysregulated form of externalization. Researchers found activations in frontal regions, in the insula, and in the striatum to be related with aggression tendencies (Dambacher F., Schuhmann T., 2015; Skibsted A.P., Cunha-Bang S., 2017; Repple J., Habel U., 2018). In another study, aggression was related with both the left superior frontal gyrus and the left middle temporal gyrus (Gong X., Quan F., 2022). Additionally, trait impulsivity that may be related with externalizing anger, was linked to prefrontal, temporal, and parietal cortices (Pan N., Wang S., 2021). From a structural point of view, a recent study used supervised machine learning to identify grey matter features related to the individual differences in externalizing anger (Consolini et al., 2022) revealing that the medial, lateral, and orbito-frontal regions, the temporal and parietal regions (temporal poles, insula, fusiform and angular gyrus, posterior cingulate), the basal ganglia, and parts of the cerebellum were found to be involved in the structural network that predicted anger externalisation.

A common neurobiological perspective views anger control as a form top-down control, involving prefrontal areas of the brain (Sorella et al., 2023). According to this view, an effort to control anger after a provocation (recreated in experimental settings with anger provocation paradigms) increases the connectivity between the amygdala and prefrontal cortices, which are responsible for top-down control (Denson et al., 2013). Accordingly, another study (Fulwiler et al., 2012) showed a positive correlation between anger control and the functional connectivity (FC) of the amygdala with the contralateral orbitofrontal cortex. Such mechanisms are also relevant for the understanding of individual differences in anger control. For example, it has been reported that violent offenders’ resting state activity after being provoked into anger showed increased amygdala-paralimbic connectivity and decreased amygdala-medial prefrontal cortex (mPFC) connectivity, suggesting that an inability of regulation inside the mPFC that can lead to reactive aggression (Siep et al., 2018). The amygdala and prefrontal cortices, which oversee top-down behaviour, become more connected when an individual makes an effort to control their anger in response to an insult (which is simulated in experimental settings using anger provocation paradigms) (Siep et al., 2018). Anger management and the functional connectivity (FC) of the amygdala with the oblique orbitofrontal cortex were found to be positively correlated (Fulwiler et al., 2012).

### Aims of the study

Thus, the first aim of the present study is to test the hypothesis that there is a negative relationship between the expression and control of anger. We predict that this relationship stands true: the more individuals can control anger, the less they externalize it. If this hypothesis is true (negative relationship between anger control and anger expression) someone can expect that the same neural circuit is involved in both externalizing and controlling anger.

The second aim of this study is to test the hypothesis that the same GM-WM circuit related to anger externalization may be related with anger control. To investigate the existence of a common neural circuitry for “anger expression” and “control”, we employed for the first time a data fusion unsupervised machine learning approach known as tIVA to decompose the brain into covarying GM-WM neural networks, and a Backward Regression approach to predict expression (STAXI Anger-outward) and anger control (STAXI Anger-control). In line with the previous evidence on these topics, we expect that an Affective network including subcortical structures such as the amygdala, the basal ganglia, as well as the Frontal network and temporal cortices, may be included in this network. We also expect the cerebellum in having a role on this network. The cerebellum has been linked to many cognitive and affective functions (Sorella et al., 2022). Finally, we expect that WM regions related to the connections between these areas should be in the same way related to anger externalization and control.

## Material and Methods

### Sample

Behavioural and structural MRI data from a cohort of 212 healthy participants (mean age of 26.06 ± 4.14 years, 131 M, 81 F), were extracted from the MPI-Leipzig Mind Brain-Body dataset available at OpenNeuro Dataset, http://openneuro.org, RRID:SCR_005031, accession number ds000221 (Babayan et al. 2020). This project was approved by the ethics committee of the University of Leipzig (Ethics Committee Approval Number: 097/15-ff). Participants were selected for the project based on their medical eligibility for magnetic resonance, excluding any psychiatric and neurological disorders. They also met the MRI safety requirements of the MPI-CBS (Mendes et al. 2019) and provided informed consent before the experimental sessions. The final sample was determined based on specific criteria, including the availability of structural images, ages falling within the range of 20 to 45 years (age categories were represented in 5-year bins, with the middle value used for calculations), and the availability of scores from STAXI.

### Questionnaire data

The State and Trait Anger Expression Inventory-2 (STAXI-2; Spielberger, 1988) was considered to assess anger facets. In line with the aim of the present study, we considered participants’ score on the subscale Anger-Expression-Out that refers to the extent to which people express their anger outwardly in a poorly controlled manner (i.e., the externalization of anger; 8 items, e.g., “I do things like slam doors”), and Angel Control subscales refer to people’s ability to monitor and control their emotions, by calming down (Anger-Control-In; 8 items, e.g., “I control my angry feelings”) and avoiding anger externalization through physical and verbal expressions (Anger-Control-Out; 8 items, e.g., “I control my behaviour”). Participants had a mean of 11.929 (SD=3.33) for anger externalization and of 22.726 (SD=3.78) for anger control.

### MRI data

The MPI-Leipzig Mind Brain-Body dataset comprises anatomical, functional, and resting-state data collected at the Day Clinic for Cognitive Neurology, University of Leipzig, utilising a 3 T Siemens Magnetom Verio scanner (Magnetom Verio, Siemens Healthcare, Erlangen, Germany) equipped with a 32-channel Siemens head coil. Each participant underwent a comprehensive imaging protocol, which included a high-resolution structural scan, four resting-state fMRI scans, two gradient echo fieldmaps, two pairs of spin echo images with reversed phase encoding direction, and a low-resolution structural image acquired using a Fluid Attenuated Inversion Recovery (FLAIR) sequence, typically employed in clinical protocols (Mendes, N., et al; 2019). For the purposes of our research, we exclusively utilised the structural images available within the MPI-Leipzig Mind Brain-Body database. These structural images were acquired using the MP2 RAGE sequence (Marques, J. P., 2010) and were characterised by the following parameters: TR = 5000 ms; TE = 2.92 ms; TI1 = 700 ms; TI2 = 2500 ms; flip angle 1 = 4°; flip angle2 = 5°; voxel size = 1.0 mm isotropic; FOV = 256 × 240 x 176 mm; bandwidth = 240 Hz/Px; GRAPPA acceleration with iPAT factor 3 (32 reference lines); pre-scan normalisation; and a total acquisition duration of 8.22 minutes.

### Preprocessing

All structural MRI (sMRI) images underwent preprocessing using SPM12 (SPM, https://www.fil.ion.ucl.ac.uk/spm/software/spm12/, RRID:SCR_007037). Initially, we conducted a comprehensive data quality check to identify and address potential distortions, including issues like head motion or artefacts. Subsequently, we executed the reorientation procedure to align the images according to a common reference point and conducted image segmentation to delineate grey matter, white matter, and cerebrospinal fluid using the CAT12 toolbox (Computational Anatomy Toolbox for SPM, http://www.neuro.uni-jena.de/cat/, RRID:SCR_019184). For the current research, only grey matter images were utilised in our analyses. To facilitate registration, we employed the Diffeomorphic Anatomical Registration using Exponential Lie algebra (DARTEL) tools for SPM12 (https://github.com/scanUCLA/spm12-dartel). Finally, we conducted image normalisation to the MNI (Montreal Neurological Institute) space, followed by spatial smoothing using a Gaussian kernel with a full width at half maximum of. (Sorella et al., 2021)

### Machine Learning analyses

To investigate the neural underpinnings of the anger constructs of interest, we employed the unsupervised machine learning method known as Transposed Independent Vector Analysis (tIVA) (Adali, Levin-Schwartz, & Calhoun, 2015; Adali, Levin-Schwartz, & Calhoun, 2015). Transposed Independent Vector Analysis (tIVA) is a blind source separation method (BSS), which provides a fully multivariate approach and enables fusion of data from multiple modalities, such as GM and WM and then decompose the brain into joint GM-WM profiles (Adali, Levin-Schwartz, & Calhoun, 2015; Adali, Levin-Schwartz, & Calhoun, 2015). The tIVA method is an extension of Independent Component Analysis (ICA) that exports statistical independence and generalizes ICA to multiple datasets by analyzing data across datasets, enabling multimodal fusion. tIVA was applied to structural data by using the Fusion ICA Toolbox (FIT, http://mialab.mrn.org/software/fit) (Calhoun, Adali, Pearlson, & Kiehl, 2006) in the MATLAB 2018a environment (https://it.mathworks.com/products/matlab.html) (*MATLAB (R2018a)*). The number of components for both modalities was estimated via the minimum norm criteria. To investigate the reliability of each modality, the ICASSO (Himberg, Hyvarinen, & Esposito, 2004; Himberg & Hyvarinen, 2003) and the Infomax algorithm were used. The resulting output consisted of a matrix with the number of subjects (rows) and the loading coefficients for each component (columns). Loading coefficients represent how each component is expressed for every participant. Subsequently, the independent componentswere translated into Talairach coordinates. Finally, the significant networks were plotted in Surf Ice (https://www.nitrc.org/projects/surfice/) (Rorden).

## Results

### Behavioural result

To explore the relationship between anger externalization and control, we performed a correlation analysis, which revealed a significant negative association between the two scales (rho = -0.418, p = 0.001). See Figure 1 A. To assess eventual effects of gender we conducted two sample t-tests to explore the potential influence of gender on anger externalization (t = 2.968; p = 0.003) and anger control (t = -4.436; p = 0.001) which confirmed gender differences. Additionally, we performed a correlation analysis to assess the impact of age on both anger externalization and control. Anger externalization was not correlated with age (rho=0.046, p=0.502) as well as anger control rho=-0.005, p=0.940). See Figure 1 B and C.

**Figure 1:**
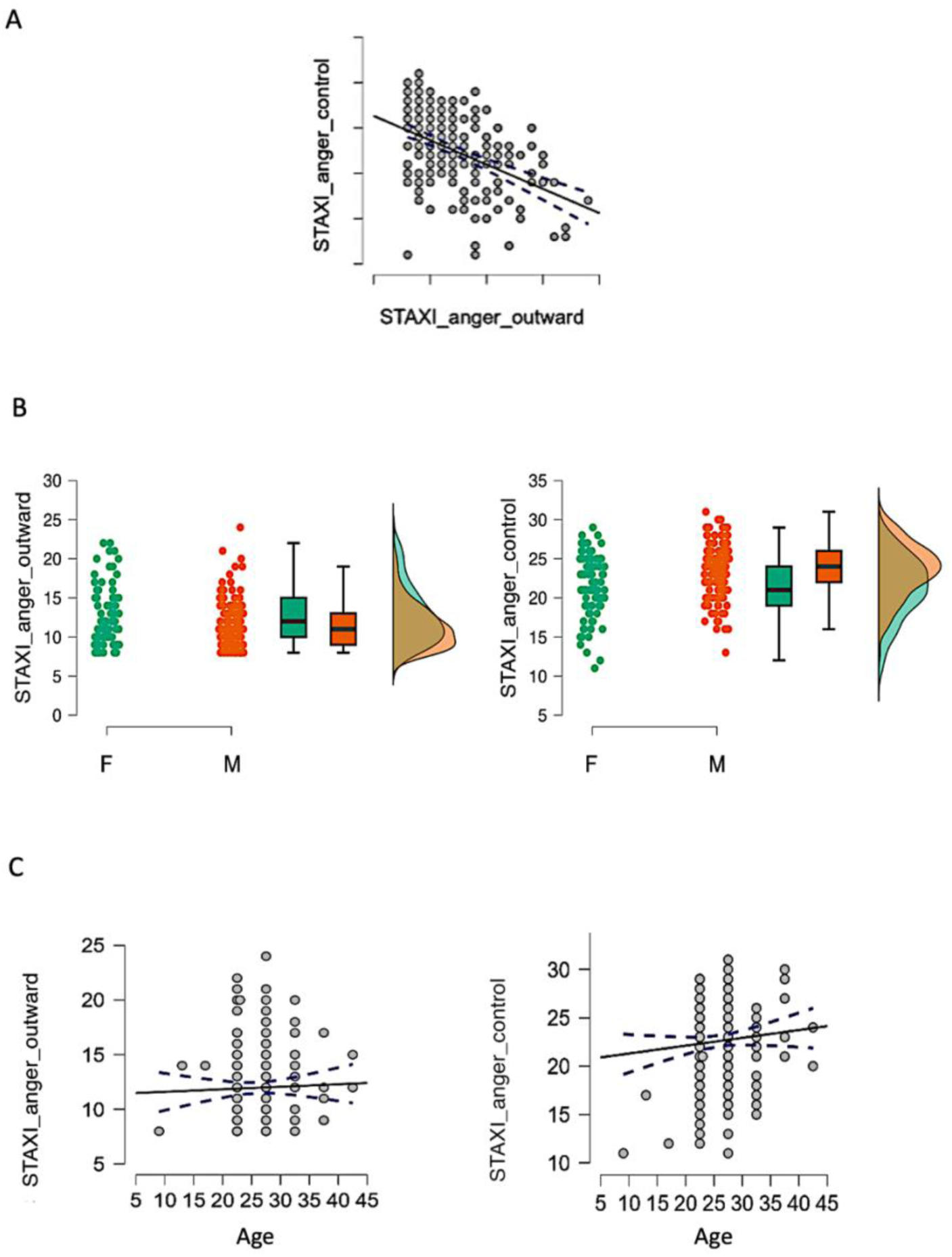
Behavioral results. A. Correlation between anger control and anger externalization. B. t-tests for gender effects in anger externalization and control. C) Correlations between anger externalization, control and age.

### Neural results

The Information Theoretic Criteria (Wax & Kailath, 1985) estimated 8 networks of covarying gray matter (GM) and white matter (WM) that were subsequently estimated via tIVA. Each component included an estimated GM component and a corresponding estimated WM component with a similar pattern of concentration across subjects. Positive values indicated increased concentration of GM/WM, while negative values a decreased concentration. The loading coefficients of these 8 components were entered two backward regression analyses, one to predict anger externalization and one to predict anger control. For anger externalization, the final model was significant (R = 0.178, R^2^ = 0.032, Adjusted R^2^ = 0.027, RMSE = 3.288, F = 6.872, p = 0.009), and included tIVA5 (beta=65.882, p = 0.009), and the intercept (beta= 1.808, p=0.641). The higher the externalization, the higher the GM-WM concentration inside this network. For anger control, the final model was significant (R = 0.205, R^2^ = 0.0042, Adjusted R^2^ = 0.037, RMSE = 3.713, F = 9.207, p = 0.003), and included again tIVA5 (beta= -86.127, p = 0.003), and the intercept (beta=35.958, p<0.001). The higher the control, the lower the GM-WM concentration inside this network. See Table 1, 2 for a description of the areas included in the network, and Figures 1, 2, for a visual representation of GM and WM as well as the residual plots. tIVA5 displayed also a significant correlation with age (rho=-0.296, p<0.001), meaning that the older the participants the lesser the GM-WM concentration. There was also a significant difference between males and females (t=3.287, p=0.001) with females showing higher GM-WM concentration compared to men.

Table 1

| **Table 1** |  |  |  |
| --- | --- | --- | --- |
| **Area** | **Brodmann**<br>**Area** | **volume**<br>**(cc)** | **random effects: Max Value (MNI, x, y, z)** |
| Paracentral<br>Lobule | 4, 5, 6, 31 | 2.2/1.9 | 6.3 (0, -35, 53)/5.7 (3, -32, 53) |
| Superior Frontal<br>Gyrus | 6, 8, 9 | 1.0/1.1 | 6.2 (0, 11, 48)/6.0 (1, 8, 51) |
| Medial Frontal<br>Gyrus | 6, 8, 9, 10,<br>32 | 2.1/2.9 | 5.4 (-1, 11, 44)/5.5 (1, 25, 42) |
| Cingulate Gyrus | 24, 32 | 1.0/1.0 | 5.0 (-1, 14, 41)/5.5 (1, 21, 40) |
| Precuneus | 7 | 0.4/0.2 | 5.2 (0, -36, 47)/4.0 (3, -34, 45) |
| Extra-Nuclear | 13 | 0.0/0.3 | -999.0 (0, 0, 0)/4.2 (43, 11, -9) |
| Superior<br>Temporal Gyrus | 22, 38 | 0.1/0.6 | 3.7 (-40, 11, -12)/4.2 (45, 8, -7) |
| Anterior<br>Cingulate | 32 | 0.1/0.3 | 4.2 (0, 36, 20)/4.0 (3, 35, 23) |
| Inferior Frontal<br>Gyrus | 47 | 0.1/0.3 | 3.6 (-37, 11, -14)/4.1 (42, 14, -12) |
| Insula | 13 | 0.1/0.3 | 3.5 (-43, 8, -5)/4.0 (45, 4, -5) |
| Sub-Gyral | * | 0.1/0.0 | 3.7 (-42, 11, -8)/-999.0 (0, 0, 0) |

Table 2

| **Table 2** |  |  |  |
| --- | --- | --- | --- |
| **Area** | **Brodmann** | **volume** | **random effects: Max Value (MNI, x, y, z)** |

|  | Area | (cc) |  |
| --- | --- | --- | --- |
| Middle Frontal Gyrus | 6, 8, 9, 10 | 1.2/1.5 | 5.6 (-27, 41, 37)/5.1 (40, 49, 20) |
| Postcentral Gyrus | 2, 3, 4, 5, 7 | 1.2/0.4 | 5.4 (-27, -43, 67)/4.7 (30, -32, 65) |
| Precentral Gyrus | 4, 6, 9 | 1.0/0.3 | 5.1 (-33, -19, 59)/4.2 (15, -27, 64) |
| Inferior Parietal Lobule | 40 | 0.4/0.6 | 4.4 (-48, -42, 39)/4.8 (52, -40, 39) |
| Sub-Gyral | * | 0.1/0.3 | 3.7 (-30, -45, 52)/4.8 (16, -25, 61) |
| Superior Frontal Gyrus | 6, 8, 9, 10 | 0.8/0.6 | 4.6 (-24, 44, 36)/4.2 (43, 49, 22) |
| Superior Parietal Lobule | 7 | 0.3/0.0 | 4.5 (-31, -45, 59)/-999.0 (0, 0, 0) |
| Uncus | 38 | 0.1/0.0 | 4.2 (-24, 13, -30)/-999.0 (0, 0, 0) |
| Supramarginal Gyrus | * | 0.2/0.1 | 4.2 (-48, -40, 35)/3.6 (53, -38, 36) |
| Precuneus | 19 | 0.2/0.0 | 4.1 (-30, -75, 40)/-999.0 (0, 0, 0) |
| Superior Temporal Gyrus | * | 0.2/0.0 | 3.9 (-24, 16, -32)/-999.0 (0, 0, 0) |
| Middle Occipital Gyrus | 18 | 0.1/0.0 | 3.8 (-31, -91, 10)/-999.0 (0, 0, 0) |
| Middle Temporal Gyrus | 21 | 0.1/0.1 | 3.7 (-40, 7, -32)/3.5 (56, -51, 5) |
| Cuneus | * | 0.1/0.0 | 3.7 (-13, -97, 9)/-999.0 (0, 0, 0) |
| Fusiform Gyrus | * | 0.1/0.0 | 3.6 (-49, -38, -22)/-999.0 (0, 0, 0) |
| Medial Frontal Gyrus | * | 0.0/0.1 | -999.0 (0, 0, 0)/3.6 (16, 57, 0) |

**Figure 1:**
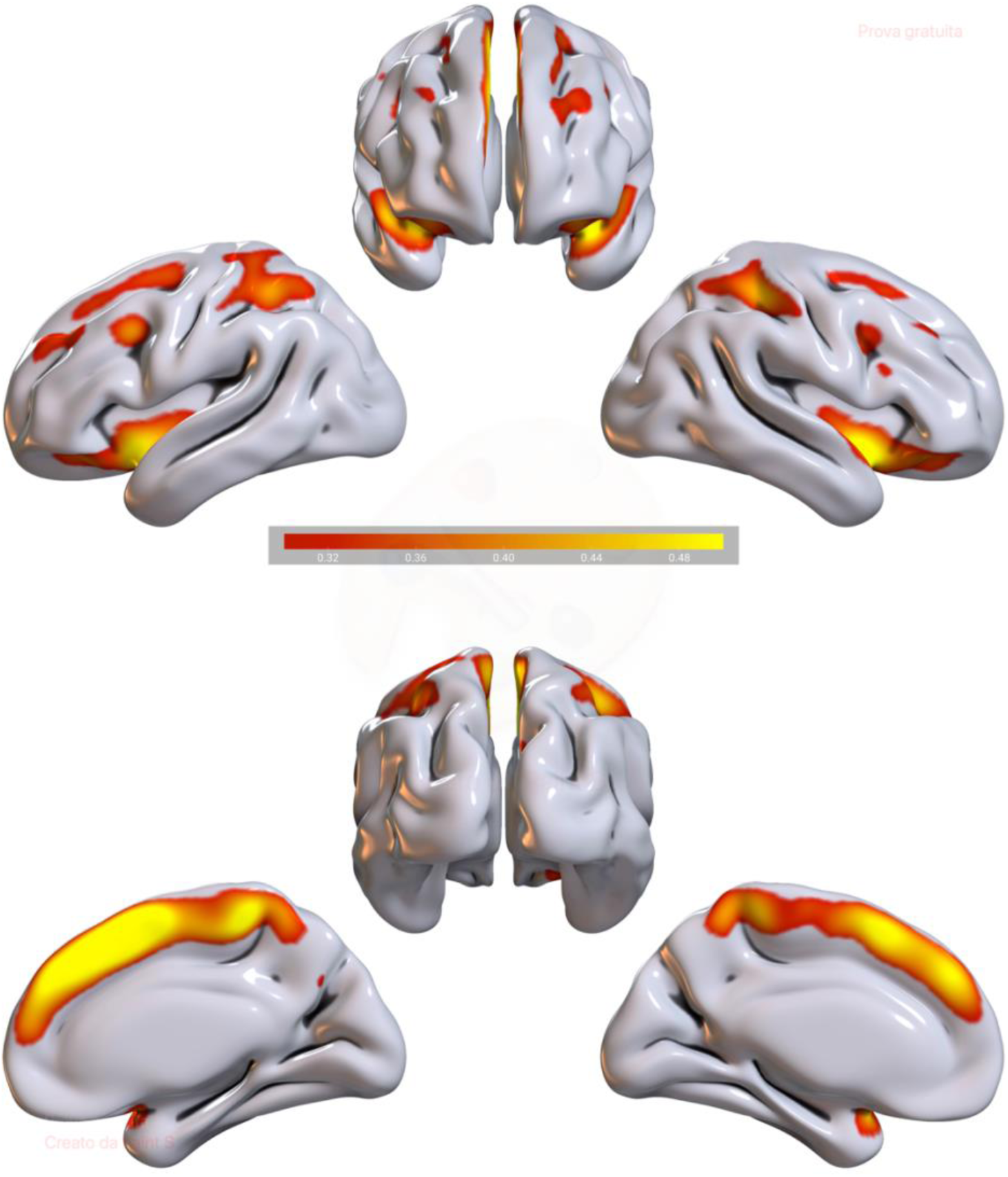
tIVA5 GM. Brain plots of tIVA5-GM. Regions with increased GM and WM are represented with warm colors. *Note*: the threshold was set at z-score > 2.

**Figure 2:**
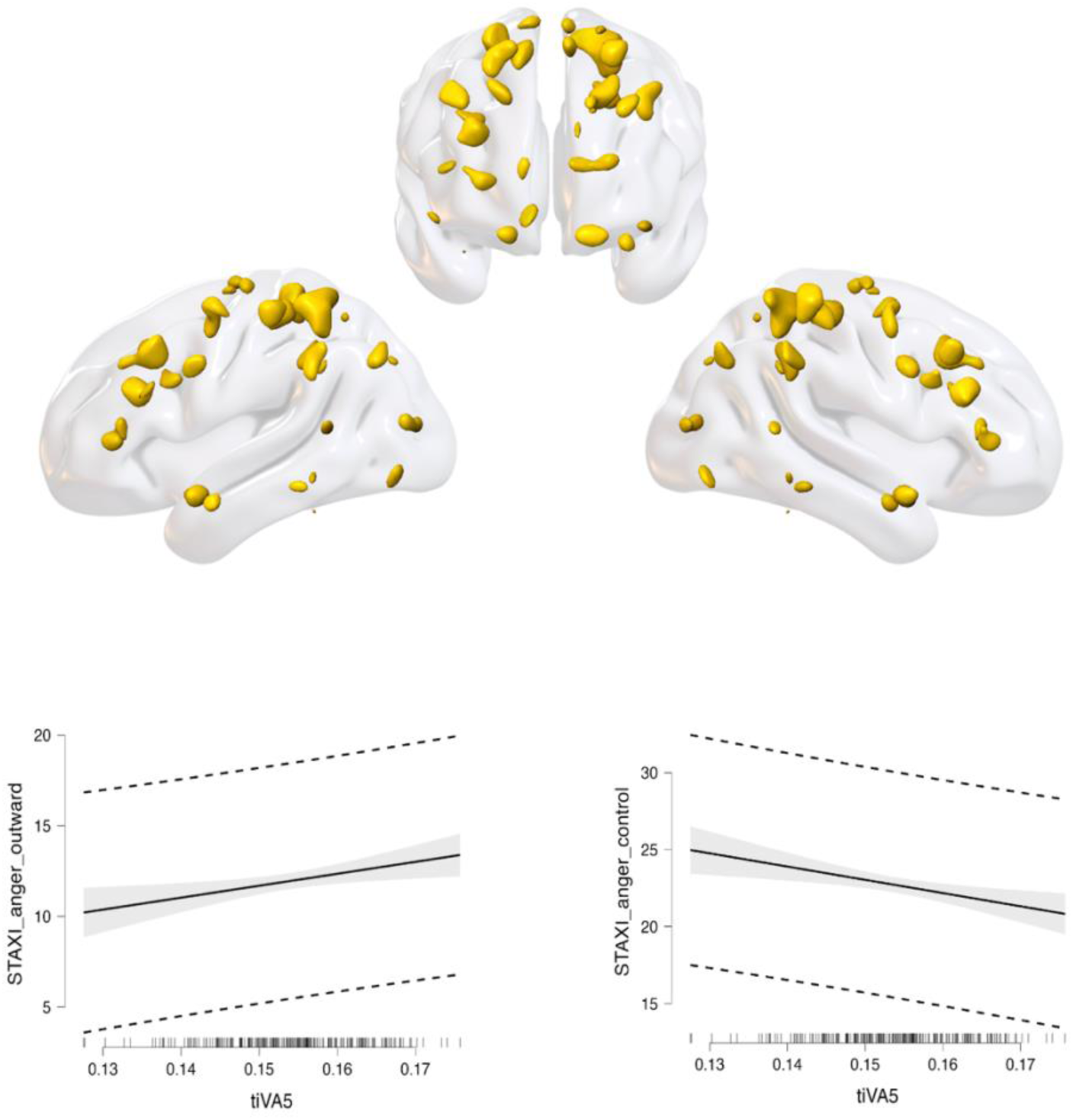
tIVA5 WM. 3D surface reconstruction plots of tIVA5-WM. *Note*: the threshold was set at z-score > 2. In the bottom part of the figure the regression residuals plots are displayed, showing that tIVA5 positively correlates with anger externalization and negatively with anger control.

## Discussion

The aim of this study was to test two hypotheses. The first hypothesis was that there was a negative relationship between anger externalization and anger control. The second hypothesis was that both the expression and control of anger could be predicted, at the neural level, by the same GM-WM network. Specifically, that a frontal control network may be involved in controlling and externalizing anger. To test these hypotheses, we took into consideration behavioural scores from the STAXI-2 subscales of anger out and anger control of 212 healthy participants, as well as their GM and WM images. We found a significant negative correlation between anger externalization and control as predicted. Moreover, by applying an unsupervised machine learning method known as Transposed Independent Vector Analysis, we found that one specific GM-WM network was able to predict both the externalization and the control of anger. Departing from previous studies, we considered both GM and WM features in the same model. This allowed us to capture comprehensive information about both aspects of brain structure, as certain psychological processes rely on both tissue types (Baggio et al., 2023). In the following paragraphs we provide details of these results.

At a behavioural level, we confirmed our hypothesis of a negative correlation between anger externalization and anger control. Thus, the individual tendency in directing anger outside (e.g. through aggressive behaviours) and the difficulty in controlling, calming down, and monitoring the outcomes of anger are often concomitant. It follows that anger externalization and control are two manifestations of a common core difficulty in regulating anger. This result provides an empirical confirmation of a clinical intuition but also aligns with the Integrative Cognitive Model (Wilkowski & Robinson, 2008; Wilkowski & Robinson, 2010). According to this model, controlling anger requires a form of effortful control, and individual differences in providing such cognitive effort can importantly influence anger externalization, mitigating aggression and reactivity in presence of more effortful control resources. This model could also explain some aspects of anger dysregulation in many psychopathologies such as borderline personality disorder (De Panfilis et al. 2019) and antisocial personality disorder (Kolla et al. 2017), characterized by this combination of high anger externalization and low anger control.

Beyond self-report data, in the present study we provided, for the first time, a neurobiological explanation of this association, with the identification of a GM-WM circuit (tIVA5) that predicted both variables, with higher concentrations predicted by higher the tendency to externalize, and lower anger control. Conceptually, one can interpret this finding as a manifestation of a possible “anger regulation continuum” between externalization and control that is reflected inside the GM-WM features of this network. Higher concentration in this anger regulation circuit corresponds on one extreme to high externalization/low control, whereas lower concentration reflects low externalization/high anger control. The placement of single individuals on this continuum can offer a useful perspective for both clinical and research reflections concerning individual anger management, for the identification of potential tailored interventions, and assessment points.

The observation of the specific brain areas predicting individual differences in anger regulation may also shed light on psychological mechanisms that influence such individual differences. The most extended component of the circuit responsible for individual differences in anger regulation was in supplemental motor areas of (SMA) in the paracentral lobule, with anterior extension towards the executive areas in medial frontal gyrus and superior frontal gyrus. The importance of emotional motor control, and motor planning have relevant implications for anger regulation (Friedman & Robbins, 2022). Consistently, the involvement of SMA may explain individual differences in anger regulation in terms of differences in threshold for action. Coherently, a previous study found that trait anger modulates the brain connectivity between the bilateral supplementary motor areas and the right frontal pole (Kim et al. 2022). According to Kohn and colleagues (2014), the SMA should be involved in execution of regulation initiated by frontal areas, also detected in the present study, and recognized in emotion regulation literature (for meta-analyses see Kohn et al., 2014; Messina et al., 2015). For example, it has been affirmed that medial prefrontal regions should play a major role in anger regulation evaluating potential outcomes and directing behaviors toward anger-eliciting stimuli (Gilam and Hendler 2015). Interestingly, a previous study coherently found that the electrical stimulation of the medial prefrontal cortex with transcranial direct current stimulation reduced anger reactions during an interpersonal game (Gilam et al. 2018). Finally, the involvement of prefrontal medial regions (including the cingulate) was found by a recent supervised machine learning study to predict individual differences in anger externalization from GM feature only (Grecucci et al. 2022).

Among prefrontal areas, also the inferior frontal gyrus has been detected in the present study. One recent meta-analysis by Sorella and colleagues (2021) showed that the right inferior frontal gyrus is active for both the perception of angry stimuli and the subjective experience of anger. This may indicate that the higher the GM concentration in this circuit the higher the anger experiences (and thus the externalization).

Another important region in our study was the Cingulate cortex. The cingulate, especially the anterior part, as well as the medial prefrontal area, have been associated to angry rumination (Denson et al. 2009), which characterizes the internalization of anger (Consolini et al. 2022). Indeed, in one study, authors found that after the presentation of angry faces, the negative connectivity of the ventral anterior cingulate cortex with the amygdala was reduced in individuals with high appetitive motivation (associated with aggression) (Passamonti et al. 2008). The involvement in the anterior cingulate in rumination and internalization of anger may appear contradictory to our findings, which show a positive association between the cingulate and externalization. However, in our study, the cingulate area was more centrally located and posterior compared to the anterior cingulate observed in previous studies on rumination and internalization (Denson et al. 2009; Consolini et al. 2022).

Another interesting region found inside the tIVA5 network was the Precuneus. Although the precuneus serves numerous functions, it is known to be the main hub of the posterior Default mode network. In a recent study it was found that anger control was associated with the default mode network (Sorella et al. 2022). The role of DMN in anger reaction/control could be responsible for hostile bias and reactivity in front of anger eliciting stimuli. In particular, the default mode network is responsible for self-referential processes and rumination that are implicated in the experience of anger.

We also found a role for the Insula. This is not surprising because the insula and other limbic and subcortical areas are all involved in the emotional experience (Picó-Pérez et al., 2018). For example, the insula has been found in angry reactions toward unfair behaviors in decision making tasks (Grecucci et al., 2013a,b). Also, according to Alia-Klein and colleagues (Alia-Klein et al. 2020), uncontrollable anger relies upon a subcortical low-road including the insula and the amygdala. In another study it has been found that in response to prohibitive language, individuals with specific genotypes, are more likely to experience anger, and that this relies on insula and right hippocampus activity (Alia-Klein et al. 2009).

For what concerns the white matter side of the tIVA5 network, results indicate a diffuse effect of white matter portions adjacent to the relative main GM regions detected by the algorithm. Several WM portions have been found close to frontal and temporal regions, thus supporting the functionality of the GM regions they communicate with. Of note, the method used in this study (tIVA) does not provide information on white matter fibers integrity as usually DTI does. WM was used in a similar way as GM was, with the idea of detecting WM density and its association with anger externalization and control (see Baggio et al., 2023; Grecucci et al., 2023; 2024; Jornkokgoud, et al., 2024 for similar methods).

Another intriguing finding of the present study was that anger externalization and control displayed were different in males and females. In our sample, females were found to have higher externalization and coherently lower control compared to men. Coherently, the anger regulation circuit presented loading coefficients were higher in women than in men (higher GM-WM concentration in females than males). This result can be somewhat counterintuitive because we know from the literature that the tendency for men to aggress is more pronounced in man than women (at least for aggression that produced pain or physical injury), whereas perceived aggressivity (e.g. perception that enacting a behaviour would produce harm to the target, guilt and anxiety in oneself) is higher in females (Eagly & Steffen, 1986; Bettemcourt & Miller, 1996). There are some possible explanations for why some women may experience higher levels of anger than men. One possibility is that women are more sensitive to social injustices, and experience gender stereotypes, injustice, role conflicts, lack of decision-making power, and even harassment or abuse, social and familial pressures (Thomas, 2005). Alternatively, it is possible that female participants in the present study are not representative of the general population. Finally, we should consider that self-reported anger expression/control does not coincide with behavioural manifestation of anger. In order to draw clear conclusions regarding gender differences in anger management and the related brain correlates, future studies should investigate such differences on the basis of observed behaviours.

## Conclusions and Limitations

This study expands on previous research on the neural bases of anger, by adding new evidence that a network of covarying GM-WM predicted anger externalization and control. These findings may help future treatment of people suffering from anger dysregulation. Several psychopathologies are associated with excessive anger externalization or lack of control, including borderline personality disorder, antisocial personality disorder, intermittent explosive disorder, and anxiety disorders (De Panfilis et al., 2019; Kolla et al., 2017; Coccaro et al., 2014). On the other hand, excessive anger control characterizes anxiety disorders and dependent personality disorder (Grecucci et al., 2020). In this context, this study reveals potential clinical value, as it could be used for diagnostic or predictive purposes regarding the onset of anger-related disorders. This circuit could be considered a potential target of neurostimulation treatment to reduce anger externalization and increase anger control.

This study does not come without limitations. A limitation concerns the exclusive use of healthy participants only. An interesting hypothesis is that people suffering from anger related problems may display even larger effects in the same brain network. Future studies could consider this investigation to enhance therapeutic approaches for such specific groups. Second, we investigated individual differences in anger regulation on the basis of self-reported regulation. Another limitation is that this study focused only on structural analysis, leaving us without knowledge of activation patterns related to anger externalization/control. Future studies may want to explore the possibility of fusing structural and functional MRI data to expand these results. While further research is necessary, we believe that these findings could provide valuable insights for understanding and in the future predicting anger related problems. The available data could ultimately assist healthcare practitioners in developing specific psychological treatments, such as targeted interventions involving brain stimulation/modulation or pharmacological techniques, for individuals experiencing anger-related issues.

## Author contribution

AG, FG, EM conceptualized the study, analyzed the data, wrote the manuscript; XY, GS, IM, wrote and reviewed the manuscript, IM, GS conceptualized the study

## References

1. Achenbach, T. M., Ivanova, M. Y., Rescorla, L. A., Turner, L. V., & Althoff, R. R. (2016). Internalizing/externalizing problems: Review and recommendations for clinical and research applications. Journal of the American Academy of Child & Adolescent Psychiatry, 55(8), 647–656.

2. Adali, T., Levin-Schwartz, Y., & Calhoun, V. D. (2015). Multimodal Data Fusion Using Source Separation: Two Effective Models Based on ICA and IVA and Their Properties. Proceedings of the IEEE, 103(9), 1478–1493. 10.1109/JPROC.2015.2461624

3. Alia-Klein N., Gan G., Gilam G., Bezek J., Bruno A., Denson T.F., Hendler T., Lowe L., Mariotti V., Muscatello M.R., Palumbo S., Pellegrini S., Pietrini P., Rizzo A., Verona E., The feeling of anger: from brain networks to linguistic expressions, Neurosci. Biobehav. Rev. 108 (2020) 480–497, 10.1016/j.neubiorev.2019.12.002

4. Archer J. (2004) Sex differences in aggression in real-world settings:a meta-analytic review. Rev Gen Psychol 8:291–322. 10.1037/1089-2680.8.4.291

5. Archer J. (2009) Does sexual selection explain human sex differences in aggression? Behav Brain Sci 32:249–266. 10.1017/S0140525X09990951

6. Babayan, A.; Blazeij Baczkowski; Cozatl, R.; Dreyer, M.; Engen, H.; Erbey, M.; Falkiewicz, M.; Farrugia, N.; Gaebler, M.; Golchert, J.; Golz, L.; Gorgolewski, K.; Haueis, P.; Huntenburg, J.; Jost, R.; Yelyzaveta Kramarenko; Krause, S.; Kumral, D.; Lauckner, M.; Margulies, D. S.; Mendes, N.; Ohrnberger, K.; Oligschläger, S.; Osoianu, A.; Pool, J.; Reichelt, J.; Reiter, A.; Röbbig, J.; Schaare, L.; Smallwood, J.; Villringer, A. MPI-Leipzig_Mind-Brain-Body, 2020. 10.18112/OPENNEURO.DS000221.V1.0.0

7. Baggio, T., Grecucci, A., Meconi, F., Messina, I. (2023). Anxious brains: A combined data fusion machine learning approach to predict trait anxiety from morphometric features. Sensors, 23,2.

8. Baron, K. G., Smith, T. W., Butner, J., Nealey-Moore, J., Hawkins, M. W., & Uchino, B. N. (2006). Hostility, anger, and marital adjustment: Concurrent and prospective associations with psychosocial vulnerability. Journal of Behavioral Medicine, 30(1), 1–10. https://link.springer.com/article/10.1007/s10865-006-9086-z

9. Bettencourt, B. A., Talley, A., Benjamin, A. J., & Valentine, J. (2006). Personality and aggressive behavior under provoking and neutral conditions: A meta-analytic review. Psychological Bulletin, 132(5), 751–777. 10.1037/0033-2909.132.5.751

10. Bettencourt, B., & Miller, N. (1996). Gender differences in aggression as a function of provocation: a meta-analysis. Psychological bulletin, 119(3), 422.

11. Bettencourt B. A., Miller N (1996) Gender differences in aggression as a function of provocation: a meta-analysis. Psychol Bull 119:422–447. 10.1037/0033-2909.119.3.422

12. Birkley, E. L., & Eckhardt, C. I. (2015). Anger, hostility, internalizing negative emotions, and intimate partner violence perpetration: A meta-analytic review. Clinical psychology review, 37, 40–56.

13. Bridewell, W. B., & Chang, E. C. (1997). Distinguishing between anxiety, depression, and hostility: Relations to anger-in, anger-out, and anger control. Personality and Individual Differences, 22(4), 587–590.

14. Carver C.S., E. Harmon-Jones (2009). Anger is an approach-related affect: evidence and implications, Psychol. Bull. 135 (2) 183–204, 10.1037/%20a0013965

15. Coccaro, E. F., Lee, R., & Mccloskey, M. S. (2014). Relationship between psychopathy, aggression, anger, impulsivity, and intermittent explosive disorder. Aggressive Behavior, 40(6), 526–536. 10.1002/ab.21536

16. Consolini, J., Sorella, S., & Grecucci, A. (2022). Evidence for lateralized functional connectivity patterns at rest related to the tendency of externalizing or internalizing anger. *Cognitive*, Affective and Behavioral Neuroscience, 22(4), 788–802. 10.3758/s13415-022-01012-0

17. Dambacher F., Schuhmann T., Lobbestael J., Arntz A., Brugman S., Sack A.T., Reducing proactive aggression through non-invasive brain stimulation, Soc. Cogn. Affect. Neurosci. 10 (10) (2015) 1303–1309, 10.1093/scan/nsv018

18. De Panfilis, C., Schito, G., Generali, I., Gozzi, L., Ossola, P., Marchesi, C., & Grecucci, A. (2019). Emotions at the border: Increased punishment behavior during fair interpersonal exchanges in borderline personality disorder. Journal of Abnormal Psychology, 128(2), 162–172. 10.1037/abn0000404

19. Denson, T. F., Ronay, R., Hippel, W. V., & Schira, M. M. (2013). Endogenous testosterone and cortisol modulate neural responses during induced anger control. Social Neuroscience, 8(2), 165–177. 10.1080/17470919.2012.655425

20. Diez I., Bonifazi P., Escudero I., Mateos B., Munoz M.A., Stramagliã S., Cortes J. M., A novel brain partition highlights the modular skeleton shared by structure and function, Sci. Rep. 5 (1) (2015) 10532, 10.1038/srep10532

21. Dodge, K. A., & Coie, J. D. (1987). Social-information-processing factors in reactive and proactive aggression in children’s peer groups. Journal of Personality and Social Psychology, 53, 1146–1158. 10.1037/0022-3514.53.6.1146

22. Eagly, A. H., & Steffen, V. J. (1986). Gender and aggressive behavior: a meta-analytic review of the social psychological literature. Psychological bulletin, 100(3), 309.

23. Farokhian, F., Yang, C., Beheshti, I., Matsuda, H., & Wu, S. (2017). Age-related gray and white matter changes in normal adult brains. Aging and Disease, 8(6), 899–909. 10.14336/AD.2017.0502

24. Feshbach, N. D., & Feshbach, S. (1969). The relationship between empathy and aggression in two age groups. Developmental Psychology, 1(2), 102–107. 10.1037/h0027016

25. Fogassi, L., & Luppino, G. (2005). Motor functions of the parietal lobe. In Current Opinion in Neurobiology (Vol. 15, Issue 6, pp. 626–631). 10.1016/j.conb.2005.10.015

26. Friedman, N. P., & Robbins, T. W. (2022). The role of prefrontal cortex in cognitive control and executive function. In Neuropsychopharmacology (Vol. 47, Issue 1, pp. 72– 89). Springer Nature. 10.1038/s41386-021-01132-0

27. Fulwiler, C. E., King, J. A., & Zhang, N. (2012). Amygdala– orbitofrontal resting-state functional connectivity is associated 14 SORELLA ET AL. with trait anger. Neuroreport, 23(10), 606–610. 10.1097/wnr.0b013e3283551cfc

28. Ghomroudi, P. A., Scaltritti, M., & Grecucci, A. (2023). Decoding reappraisal and suppression from neural circuits: A combined supervised and unsupervised machine learning approach. Cognitive, affective & behavioral neuroscience, 10.3758/s13415-023-01076-6. Advance online publication.

29. Gilam G., Hendler T., Deconstructing anger in the human brain. In social behavior from rodents to humans, in: M. Wohr, S. Krach (Eds.), Current Topics in Behavioral Neurosciences Vol. 30, Springer International Publishing, Cham, 2015, pp. 257–273, 10.1007/7854_2015_408

30. Gong X., Quan F., Wang L., Zhu W., Lin D., Xia L.-X., The relationship among regional gray matter volume in the brain, machiavellianism and social aggression in emerging adulthood: a voxel-based morphometric study, Curr. Psychol. (2022), 10.1007/s12144-022-03574-1

31. Grecucci, A., Dadomo, H., Salvato, G., Lapomarda, G., Sorella, S., Messina, I. (2023). Abnonrmal Brain Circuits characterize Borderline Personality and Mediate the Relationship Between Specific Childhood Traumas and Symptoms. A mCCA+ jICA and Random Forest Approach. Sensors, 23,5.

32. Grecucci, A., Giorgetta, C., Brambilla, P., Zanon, S., Perini, L., Balestrieri, M., Bonini, N., & Sanfey, A. (2013). Anxious ultimatums. How anxiety affects socio-economic decisions. Cognition & Emotion., 27(2), 230–244. 10.1080/02699931.2012.698982

33. Grecucci, A., Monachesi, B., Messina, I. (2024). Reduced GM-WM concentration inside the Default Mode Network in individuals with high emotional intelligence and low anxiety: a data fusion mCCA+jICA approach. Social Cognitive and Affective Neuroscience.

34. Grecucci, A., Sorella, S., & Consolini, J. (n.d.). Decoding Individual Differences in Expressing and Suppressing Anger from Structural Brain Networks: a Supervised Machine Learning Approach.

35. Grecucci, A., Sorella, S., & Consolini, J. (2023). Decoding individual differences in expressing and suppressing anger from structural brain networks: A supervised machine learning approach. Behavioural Brain Research, 439. 10.1016/j.bbr.2022.114245

36. Kolla, N. J., Meyer, J. H., Bagby, R. M., & Brijmohan, A. (2017). Trait Anger, Physical Aggression, and Violent Offending in Antisocial and Borderline Personality Disorders. Journal of Forensic Sciences, 62(1), 137–141. 10.1111/1556-4029.13234

37. Harmon-Jones E, Gable PA. (2018). On the role of asymmetric frontal cortical activity in approach and withdrawal motivation: An updated review of the evidence. Psychophysiology.;55(1). https://pubmed.ncbi.nlm.nih.gov/28459501/

38. Kelley, N. J., Hortensius, R., Schutter, D. J. L. G., & Harmon-Jones, E. (2017). The relationship of approach/avoidance motivation and asymmetric frontal cortical activity: A review of studies manipulating frontal asymmetry. International Journal of Psychophysiology, 119, 19–30. 10.1016/j.ijpsycho.2017.03.001

39. Kernberg, O. F. (2012). The inseparable nature of love and aggression: Clinical and theoretical perspectives. American Psychiatric Pub

40. Kohn, N., Eickhoff, S. B., Scheller, M., Laird, A. R., Fox, P. T., & Habel, U. (2014). Neural network of cognitive emotion regulation—an ALE meta-analysis and MACM analysis. Neuroimage, 87, 345–355

41. Kolla, N. J., Meyer, J. H., Bagby, R. M., & Brijmohan, A. (2016). Trait anger, physical aggression, and violent offending in antisocial and borderline personality disorders. Journal of Forensic Sciences, 62(1), 137–141. 10.1111/1556-4029.13234

42. Jornkokgoud, K., Baggio, T., Bakiaj, R., Wongupparaj, P., Job, R., Grecucci, A. (2024). Narcissus reflected: grey and White matter features joint contribution to the default mode network in predicting narcissistic personality traits. European journal of neuroscience.

43. Lagerspetz, Kirsti & Björkqvist, Kaj & Peltonen, Tarja. (1988). Is Indirect Aggression Typical of Females? Gender Differences in Aggressiveness in 11- to 12-Year-Old Children. Aggressive Behavior. 14. 403–414. 10.1002/1098-2337(1988)14:6%3C403::AID-AB2480140602%3E3.0.CO;2-D

44. Litt, M. D., Cooney, N. L., & Morse, P. (2000). Reactivity to alcoholrelated stimuli in the laboratory and in the field: Predictors of craving in treated alcoholics. Addiction, 95(6), 889–900. 10.1046/j.1360-0443.2000.9568896.x

45. Manfredi, P., & Taglietti, C. (2022). A psychodynamic contribution to the understanding of anger-The importance of diagnosis before treatment. Research in Psychotherapy: Psychopathology, Process, and Outcome, 25(2).

46. Marques, J. P., Kober, T., Krueger, G., van der Zwaag, W., Van de Moortele, P. F., & Gruetter, R. (2010). MP2RAGE, a self bias-field corrected sequence for improved segmentation and T1-mapping at high field. NeuroImage, 49(2), 1271–1281. 10.1016/j.neuroimage.2009.10.002

47. Mattevi, A., Sorella, S., Vellani, V., Job, R., & Grecucci, A. (2019). Regolare la rabbia: Quale strategia? Uno studio preliminare. Giornale italiano di Psicologia.

48. Mendes, N., Oligschläger, S., Lauckner, M. E., Golchert, J., Huntenburg, J. M., Falkiewicz, M., Ellamil, M., Krause, S., Baczkowski, B. M., Cozatl, R., Osoianu, A., Kumral, D., Pool, J., Golz, L., Dreyer, M., Haueis, P., Jost, R., Kramarenko, Y., Engen, H., … Margulies, D. S. (2019). Data descriptor: A functional connectome phenotyping dataset including cognitive state and personality measures. Scientific Data, 6. 10.1038/sdata.2018.307

49. Menon V., Large-scale brain networks and psychopathology: a unifying triple network model, Trends Cogn. Sci. 15 (10) (2011) 483–506, 10.1016/j.tics.2011.08.003

50. Meier J., Tewarie P., Hillebrand A., Douw L., van Dijk B.W., Stufflebeam S.M., Van Mieghem P., A mapping between structural and functional brain networks, Brain Connect 6 (4) (2016) 298–311, 10.1089/brain.2015.0408

51. Messina, I., Bianco, S., Sambin, M., & Viviani, R. (2015). Executive and semantic processes in reappraisal of negative stimuli: insights from a meta-analysis of neuroimaging studies. Frontiers in psychology, 6, 145523.

52. Ochsner, K.N.; Gross, J.J. The Neural Bases of Emotion and Emotion Regulation: A Valuation Perspective. In Handbook of emotion regulation; The Guilford Press; pp 23–42.

53. Pan N., Wang S., Zhao Y., Lai H., Qin K., Li J., Biswal B.B., Sweeney J.A., Gong Q., Brain gray matter structures associated with trait impulsivity: a systematic review and voxel-based meta-analysis, Hum. Brain Mapp. 42 (7) (2021) 2214–2235, 10.1002/hbm.25361

54. Pascual-Leone, A., Gilles, P., Singh, T., & Andreescu, C. A. (2013). Problem anger in psychotherapy: An emotion-focused perspective on hate, rage, and rejecting anger. Journal of Contemporary Psychotherapy, 43, 83–92.

55. Picó-Pérez, M., Alonso, P., Contreras-Rodríguez, O., Martínez-Zalacaín, I., López-Solà, C., Jiménez-Murcia, S., Verdejo-García, A., Menchón, J. M., & Soriano-Mas, C. (2018). Dispositional use of emotion regulation strategies and resting-state cortico-limbic functional connectivity. Brain Imaging and Behavior, 12(4), 1022–1031. 10.1007/s11682-017-9762-3

56. Repple J., Habel U., Wagels L., Pawliczek C.M., Schneider F., Kohn N., Sex differences in the neural correlates of aggression, Brain Struct. Funct. 223 (9) (2018) 4115–4124, 10.1007/s00429-018-1739-5

57. Roberton, T., Daffern, M., & Bucks, R. S. (2015). Beyond anger control: Difficulty attending to emotions also predicts aggression in offenders. Psychology of Violence, 5(1), 74.

58. Romero-Martínez A., Gonzalez M., Lilaá M., Gracia E., Martí-Bonmatí L., Álberich-Bayarri, R. Maldonado-Puig A., Ten-Esteve A., Moya-Albiol L., The brain resting-state functional connectivity underlying violence proneness: is it a reliable marker for neurocriminology? A systematic review, Behav. Sci. 9 (1) (2019) 11, 10.3390/bs9010011

59. Schieman, S. (1999). Age and Anger. In Journal of Health and Social Behavior (Vol. 40, Issue 3).

60. Skibsted A.P., Cunha-Bang S., da; Carŕe J.M., Hansen A.E., Beliveau V., Knudsen G.M., Fisher P.M., Aggression-related brain function assessed with the point subtraction aggression paradigm in FMRI, Aggress. Behav. 43 (6) (2017) 601–610, 10.1002/ab.21718

61. Sinaeifar Z., Mayeli M., Shafie M., Pooyan A., Cattarinussi G., Aarabi M.H., Sambataro F., Trait anger representation in microstructural white matter tracts: a diffusion MRI study, J. Affect. Disord. 322 (2023) 249–257, 10.1016/j.jad.2022.11.020

62. Smith, T. W., Glazer, K., Ruiz, J. M., & Gallo, L. C. (2004). Hostility, anger, aggressiveness, and coronary heart disease: An interpersonal perspective on personality, emotion, and health. Journal of Personality, 72, 1217–1270. https://onlinelibrary.wiley.com/doi/10.1111/j.1467-6494.2004.00296.x

63. Sorella, S., & Grecucci, A. (2022). THE NEURAL BASES OF ANGER. In C. R. Martin, V. R. Preedy, & V. B. Patel (Eds.), Handbook of Anger, Aggression, and Violence. Springer Cham.

64. Sorella, S., Grecucci, A., Piretti, L., & Job, R. (2021). Do anger perception and the experience of anger share common neural mechanisms? Coordinate-based meta-analytic evidence of similar and different mechanisms from functional neuroimaging studies. NeuroImage, 230. 10.1016/j.neuroimage.2021.117777

65. Sorella, S., Vellani, V., Siugzdaite, R., Feraco, P., & Grecucci, A. (2022). Structural and functional brain networks of individual differences in trait anger and anger control: An unsupervised machine learning study. European Journal of Neuroscience, 55(2), 510–527. 10.1111/ejn.15537

66. Spalletta, G., Piras, F., & Gili, T. (n.d.). *Brain Morphometry*. http://www.springer.com/series/7657

67. Spielberger, C. D. (1988). Manual for the State-Trait Anger Expression Inventory (STAXI) Psychological Assessment Resources: Odessa.

68. Spielberger, C. D., Foreyt, J. P., Goodrick, G., & Reheiser, E. C. (1995). Personality characteristics of users of smokeless tobacco compared with cigarette smokers and non-users of SORELLA ET AL. 17 tobacco products. Personality and Individual Differences, 19(4), 439–448. 10.1016/0191-8869(95)00084-j

69. Thomas, S. P. (2005). Women’s anger, aggression, and violence. Health Care for Women International, 26(6), 504–522. 10.1080/07399330590962636

70. Suarez L.E., Markelló R.D., Betzel R.F., Misic B., Linking structure and function in macroscale brain networks, Trends Cogn. Sci. 24 (4) (2020) 302–315, 10.1016/j.tics.2020.01.008

71. Vanasse T.J., Fox P.T., Fox P.M., Cauda F., Costa T., Smith S.M., Eickhoff S.B., Lancaster J. L., Brain pathology recapitulates physiology: a network meta-analysis, Commun. Biol. 4 (1) (2021) 301, 10.1038/s42003-021-01832-9

72. Wilkowski, B. M., & Robinson, M. D. (2008). The cognitive basis of trait anger and reactive aggression: An integrative analysis. Personality and social psychology review, 12(1), 3–21.

